# Virological characteristics of the SARS-CoV-2 KP.2 variant

**DOI:** 10.1101/2024.04.24.590786

**Authors:** Yu Kaku, Keiya Uriu, Yusuke Kosugi, Kaho Okumura, Daichi Yamasoba, Yoshifumi Uwamino, Jin Kuramochi, Kenji Sadamasu, Kazuhisa Yoshimura, Hiroyuki Asakura, Mami Nagashima, The Genotype to Phenotype Japan (G2P-Japan) Consortium, Jumpei Ito, Kei Sato

## Abstract

The JN.1 variant (BA.2.86.1.1), arising from BA.2.86(.1) with the S:L455S substitution, exhibited increased fitness and outcompeted the previous dominant XBB lineage by the biggening of 2024. JN.1 subsequently diversified, leading to the emergence of descendants with spike (S) protein substitutions such as S:R346T and S:F456L. Particularly, the KP.2 (JN.1.11.1.2) variant, a descendant of JN.1 bearing both S:R346T and S:F456L, is rapidly spreading in multiple regions as of April 2024. Here, we investigated the virological properties of KP.2. KP.2 has three substitutions in the S protein including the two above and additional one substitution in non-S protein compared with JN.1. We estimated the relative effective reproduction number (R_e_) of KP.2 based on the genome surveillance data from the USA, United Kingdom, and Canada where >30 sequences of KP.2 has been reported, using a Bayesian multinomial logistic model. The R_e_ of KP.2 is 1.22-, 1.32-, and 1.26-times higher than that of JN.1 in USA, United Kingdom, and Canada, respectively. These results suggest that KP.2 has higher viral fitness and potentially becomes the predominant lineage worldwide. Indeed, as of the beginning of April 2024, the estimated variant frequency of KP.2 has already reached 20% in United Kingdom.

The pseudovirus assay showed that the infectivity of KP.2 is significantly (10.5-fold) lower than that of JN.1. We then performed a neutralization assay using monovalent XBB.1.5 vaccine sera and breakthrough infection (BTI) sera with XBB.1.5, EG.5, HK.3 and JN.1 infections. In all cases, the 50% neutralization titer (NT_50_) against KP.2 was significantly lower than that against JN.1. Particularly, KP.2 shows the most significant resistance to the sera of monovalent XBB.1.5 vaccinee without infection (3.1-fold) as well as those who with infection (1.8-fold). Altogether, these results suggest that the increased immune resistance ability of KP.2 partially contributes to the higher R_e_ more than previous variants including JN.1.

## Text

The JN.1 variant (BA.2.86.1.1), arising from BA.2.86(.1) with the S:L455S substitution, exhibited increased fitness and outcompeted the previous dominant XBB lineage by the biggening of 2024.^1^ JN.1 subsequently diversified, leading to the emergence of descendants with spike (S) protein substitutions such as S:R346T and S:F456L. Particularly, the KP.2 (JN.1.11.1.2) variant, a descendant of JN.1 bearing both S:R346T and S:F456L, is rapidly spreading in multiple regions as of April 2024. Here, we investigated the virological properties of KP.2. KP.2 has three substitutions in the S protein including the two above and additional one substitution in non-S protein compared with JN.1 (**Figure 1A**). We estimated the relative effective reproduction number (R_e_) of KP.2 based on the genome surveillance data from the USA, United Kingdom, and Canada where >30 sequences of KP.2 has been reported, using a Bayesian multinomial logistic model (**Figures 1B and 1C; Table S3**).^2^ The R_e_ of KP.2 is 1.22-, 1.32-, and 1.26-times higher than that of JN.1 in USA, United Kingdom, and Canada, respectively (**Figure 1B**). These results suggest that KP.2 has higher viral fitness and potentially becomes the predominant lineage worldwide. Indeed, as of the beginning of April 2024, the estimated variant frequency of KP.2 has already reached 20% in United Kingdom (**Figure 1C**).

**Figure 1.**
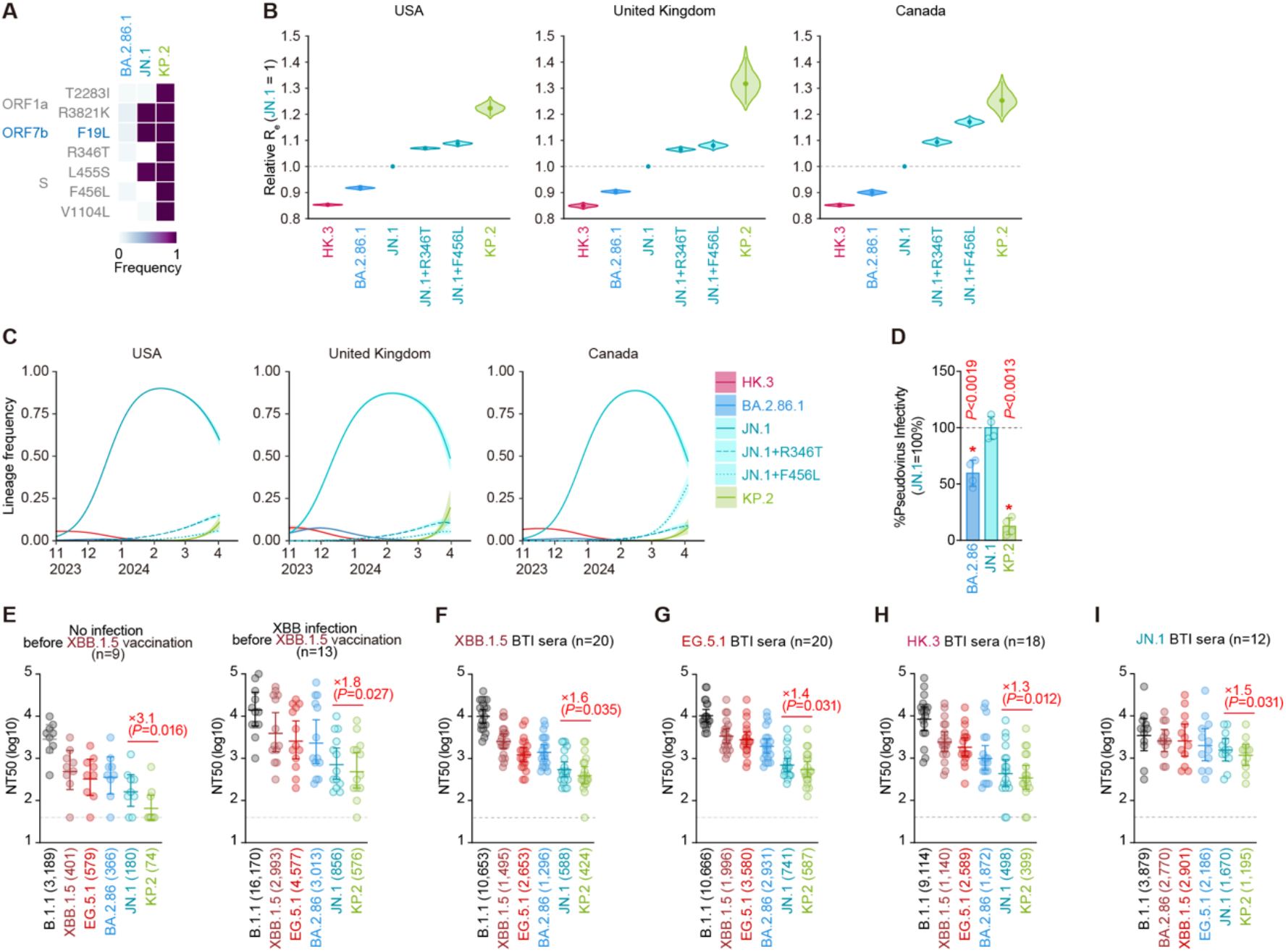
Virological features of KP.2. **(A)**Frequency of mutations in KP.2 and other lineages of interest. Only mutations with a frequency >0.5 in at least one but not all the representative lineages are shown. **(B)**Estimated relative R_e_ of the variants of interest in the USA, United Kingdom, and Canada. The relative R_e_ of JN.1 is set to 1 (horizontal dashed line). Violin, posterior distribution; dot, posterior mean; line, 95% Bayesian confidence interval. **(C)**Estimated epidemic dynamics of the variants of interest in the USA, United Kingdom, and Canada from November 1, 2023 to April 11, 2024. Countries are ordered according to the number of detected sequences of KP.2 from high to low. Line, posterior mean, ribbon, 95% Bayesian confidence interval. **(D)**Lentivirus-based pseudovirus assay. HOS-ACE2/TMPRSS2 cells were infected with pseudoviruses bearing each S protein of BA.2.86, JN.1 and KP.2. The amount of input virus was normalized to the amount of HIV-1 p24 capsid protein. The percentage infectivity of BA.2.86 and KP.2 are compared to that of JN.1. The horizontal dash line indicates the mean value of the percentage infectivity of JN.1. Assays were performed in quadruplicate, and a representative result of four independent assays is shown. The presented data are expressed as the average ± SD. Each dot indicates the result of an individual replicate. Statistically significant differences versus JN.1 is determined by two-sided Student’s *t* tests (*, *P*<0.01). (**E**-**I**) Neutralization assay. Assays were performed with pseudoviruses harboring the S proteins of B.1.1, XBB.1.5, EG.5.1, BA.2.86, JN.1 and KP.2. The following sera were used: vaccinated sera from fully vaccinated individuals who had not been infected (9 donors) and vaccinated sera from fully vaccinated individuals who had been infected with XBB subvariants (after June, 2023) (13 donors) (**E**); convalescent sera from fully vaccinated individuals who had been infected with XBB.1.5 (one 2-dose vaccinated donor, seven 3-dose vaccinated donors, six 4-dose vaccinated donors, five 5-dose vaccinated donors and one 6-dose vaccinated donor. 20 donors in total) (**F**); EG.5.1 (one 2-dose vaccinated donor, six 3-dose vaccinated donors, five 4-dose vaccinated donors, four 5-dose vaccinated donors and four 6-dose vaccinated donors. 20 donors in total) (**G**); individuals who had been infected with HK.3 (three 2-dose vaccinated donors, five 3-dose vaccinated donor, two 4-dose vaccinated donors, three 5-dose vaccinated donors, one 6-dose vaccinated donor and four donors with unknown vaccine history. 18 donors in total) (**H**) and individuals who had been infected with JN.1 (one 2-dose vaccinated donor, two 3-dose vaccinated donors, two 7-dose vaccinated donors and seven donors with unknown vaccine history. 12 donors in total) (**I**). Assays for each serum sample were performed in quadruplicate to determine the 50% neutralization titer (NT_50_). Each dot represents one NT_50_ value, and the geometric mean and 95% confidence interval are shown. The number in parenthesis indicates the geometric mean of NT_50_ values. The horizontal dash line indicates the detection limit (40-fold). Statistically significant differences versus KP.2 were determined by two-sided Wilcoxon signed-rank tests, and p values are indicated in parentheses. The fold changes of NT_50_ from that of KP.2 are indicated with “X”.

The pseudovirus assay showed that the infectivity of KP.2 is significantly (10.5-fold) lower than that of JN.1 (**Figure 1D**). We then performed a neutralization assay using monovalent XBB.1.5 vaccine sera and breakthrough infection (BTI) sera with XBB.1.5, EG.5, HK.3 and JN.1 infections. In all cases, the 50% neutralization titer (NT_50_) against KP.2 was significantly lower than that against JN.1 (**Figures 1E-I**). Particularly, KP.2 shows the most significant resistance to the sera of monovalent XBB.1.5 vaccinee without infection (3.1-fold) as well as those who with infection (1.8-fold) (**Figures 1E**). Altogether, these results suggest that the increased immune resistance ability of KP.2 partially contributes to the higher R_e_ more than previous variants including JN.1.

### Grants

Supported in part by AMED ASPIRE Program (JP24jf0126002, to G2P-Japan Consortium and Kei Sato); AMED SCARDA Japan Initiative for World-leading Vaccine Research and Development Centers “UTOPIA” (JP243fa627001h0003, to Kei Sato); AMED SCARDA Program on R&D of new generation vaccine including new modality application (JP243fa727002h0003, to Kei Sato); AMED Research Program on Emerging and Re-emerging Infectious Diseases (JP243fa727002, to Kei Sato); JST PRESTO (JPMJPR22R1, to Jumpei Ito); JSPS KAKENHI Grant-in-Aid for Early-Career Scientists (JP23K14526, to Jumpei Ito); JSPS KAKENHI Grant-in-Aid for Scientific Research A (JP24H00607, to Kei Sato); JSPS Research Fellow DC2 (JP22J11578, to Keiya Uriu); JSPS Research Fellow DC1 (JP23KJ0710, to Yusuke Kosugi); The Tokyo Biochemical Research Foundation (to Kei Sato); The Mitsubishi Foundation (to Kei Sato).

