## Supplementary Appendix for "Virological characteristics of the SARS-CoV-2 KP.2 variant"

**Table of Contents**

**Contents** Page


| Contents | Page |
|---|---|
| **Materials and Methods** | 2-5 |
| Ethics statement | |
| Human serum collection | |
| Epidemic dynamics analysis and mutation frequency calculation | |
| Plasmid construction | |
| Cell culture | |
| Pseudovirus preparation | |
| Neutralization assay | |
| Data availability | |
| **Table S1.** Human infection sera used in this study | 6 |
| **Table S2.** Human XBB.1.5 vaccine sera used in this study | 7 |
| **Table S3.** Estimated relative Re and epidemic dynamics modeling parameters of the representative SARS-CoV-2 Omicron sublineages spreading in USA, United Kingdom, and Canada from November 1, 2023 to April 11, 2024 | 8-13 |
| **Table S4.** Primers used in this study | 14 |
| **Figure S1.** Virological features of KP.2 | 15-16 |
| **Consortia** | 17 |
| **Acknowledgments** | 18 |
| **Supplemental References** | 19 |

### Materials and Methods

#### Ethics statement

All protocols involving specimens from human subjects recruited at Kyoto University, Interpark Kuramochi Clinic, Namikibashi Clinic, Wakaba Clinic and Keio University were reviewed and approved by the Institutional Review Boards of The Institute of Medical Science, The University of Tokyo (approval IDs: 2021-1-0416 and 2022-29-0915), Kyoto University (approval ID: G1309), Interpark Kuramochi Clinic (approval ID: G2021-004), and Keio University (approval ID: 20200059), respectively. All human subjects provided written informed consent. All protocols for the use of human specimens were reviewed and approved by the Institutional Review Boards of The Institute of Medical Science, The University of Tokyo (approval IDs: 2021-1-0416, 2021-18-0617 and 2022-29-0915).

#### Human serum collection

Convalescent sera were collected from fully vaccinated individuals who had been infected with XBB.1.5 (one 2-dose vaccinated, seven 3-dose vaccinated, six 4-dose vaccinated, five 5-dose vaccinated and one 6-dose vaccinated; time interval between the last vaccination and infection, 44-691 days；14-46 days after testing. n=20 in total; average age: 44.9 years, range: 15-74 years, 25.0% male) and EG.5.1 (one 2-dose vaccinated, six 3-dose vaccinated, five 4-dose vaccinated, four 5-dose vaccinated and four 6-dose vaccinated; time interval between the last vaccination and infection, 58–500 days；4–27 days after testing. n=20 in total; average age: 53.8 years, range: 27–77 years, 50% male), individuals who had been infected with HK.3 (three 2-dose vaccinated, five 3-dose vaccinated, two 4-dose vaccinated, three 5-dose vaccinated, one 6-dose vaccinated and four unknown vaccine history; time interval between the last vaccination and infection, 44-888 days；17-112 days after testing. n=18 in total; average age: 57.4 years, range: 10-84 years, 77.8% male) and individuals who had been infected with JN.1 (one 2-dose vaccinated, two 3-dose vaccinated, two 7-dose vaccinated and seven unknown vaccine history; time interval between the last vaccination and infection, 58-498 days；13-46 days after testing. n=12 in total; average age: 69.3 years, range: 31-94 years, 41.7% male). XBB.1.5 monovalent vaccine sera from fully vaccinated individuals who had not been infected (nine donors. Average age, 66.2; range, 51–89; 44.4% male) and those from fully vaccinated individuals who had been infected with XBB subvariants (thirteen donors. Average age, 49.3; range, 32–61; 69.2% male) were collected before vaccination and three-four weeks (20–29 days) after vaccination. The SARS-CoV-2 variants were identified as previously described. [1–4] Sera were inactivated at 56°C for 30 minutes and stored at –80°C until use. The details of the convalescent sera are summarized in **Table S1 and Table S2**.

#### Epidemic dynamics analysis and mutation frequency calculation

In this study, we analyzed the viral genomic surveillance data deposited in the GISAID database (https://www.gisaid.org; downloaded on April 18, 2024). We used the data of SARS-CoV-2 collected from November 1, 2023 to April 11, 2024 in this analysis. We excluded the data of SARS-CoV-2 that i) lacks collection date and PANGO lineage information; ii) was retrieved from non-human animals; iii) was sampled by quarantine; iv) was sampled from the original passage; and v) whose genomic sequence is not longer than 28,000 base pairs and contains >2% of unknown (N) nucleotide sequences. Additionally, JN.1 data except KP.2 was classified into three groups and renamed respectively: JN.1 (JN.1 lineages without the S:R346T and S:F456L substitutions), JN.1+R346T (JN.1 lineages with S:R346T but not S:F456L), and JN.1+F456L (JN.1 lineages with S:F456L but not S:R346T). JN.1 lineages with S:R346T and S:F456L substitutions but not named KP.2 were excluded from analysis. In the downstream analysis, we only used sequences for PANGO lineages with >20 sequences in each country in the dataset. We modeled the epidemic dynamics of variants of interest in USA, Japan, and United Kingdom, where >30 genomic sequences of KP.2 were detected. The daily frequency of each viral lineage was counted. Then, epidemic dynamics and $R_e$ value for each viral were subsequently estimated according to the Bayesian multinomial logistic model, described in our previous study.[4] Briefly, we estimated the logistic slope parameter $\beta_l$ for each lineage and then calculated a relative $R_e$ for each lineage ($r_l$) as $r_l = \exp(\gamma\beta_l)$ where $\gamma$ is the average viral generation time (2.1 days) (http://sonorouschocolate.com/covid19/index.php?title=Estimating_Generation_Time_Of_Omicron). For parameter estimation, the intercept and slope parameters of JN.1 were fixed at 0. The relative $R_e$ of JN.1 was fixed at 1, and that of other lineage was estimated with respect to that of JN.1. Parameter estimation was performed by using the Markov chain Monte Carlo (MCMC) approach implemented in CmdStan v2.34.1 (https://mc-stan.org) accessed through the CmdStanr v0.6.1 R interface (https://mc-stan.org/cmdstanr/). Four independent 5,000-step MCMC chains were run including 1,000-step warmup iterations. We confirmed that an estimated $\hat{R}$ convergence diagnostic value is <1.05 and bulk and tail effective sampling sizes are >200, indicating that all runs were successfully convergent. Information on the estimated parameters is summarized in **Table S3**. Mutation frequency of each lineage was calculated by dividing the number of sequences harboring the substitution of interest with the total number of sequences in each lineage.

**Plasmid construction**

Plasmids expressing the SARS-CoV-2 spike proteins of B.1.1, BA.2.86, XBB.1.5, EG.5.1 and JN.1 were prepared in our previous studies.[5–8] Plasmids expressing the spike protein of KP.2 was generated by site-directed overlap extension PCR using pC-SARS2-S JN.1 as the template and the primers listed in **Table S4**. The resulting PCR fragment was subcloned into the KpnI-NotI site of the pCAGGS vector[9] using In-Fusion HD Cloning Kit (Takara, Cat# Z9650N). Nucleotide sequences were determined by DNA sequencing services (Eurofins), and the sequence data were analyzed by SnapGene software v6.1.1 (www.snapgene.com).

**Cell culture**

The Lenti-X 293T cells (Takara, Cat# 632180) and HOS-ACE2/TMPRSS2 cells (kindly provided by Dr. Kenzo Tokunaga), a derivative of HOS cells (a human osteosarcoma cell line; ATCC CRL-1543) stably expressing human ACE2 and TMPRSS2,[10,11] were maintained in Dulbecco's modified Eagle's medium (DMEM) (high glucose) (Wako, Cat# 044- 29765) containing 10% fetal bovine serum (Sigma-Aldrich Cat# 172012-500ML), 100 units penicillin and 100 ug/ml streptomycin (Sigma-Aldrich, Cat# P4333-100ML).

**Pseudovirus preparation**

Pseudoviruses were prepared as previously described.[5,7,8,12] Briefly, lentivirus (HIV-1)-based, luciferase-expressing reporter viruses were pseudotyped with the SARS-CoV-2 S. One prior day of transfection, the LentiX-293T cells were seeded at a density of $2 \times 10^6$ cells. The LentiX-293T cells were cotransfected with 1 μg psPAX2-IN/HiBiT (a packaging plasmid encoding the HiBiT-tag-fused integrase[10]), 1 μg pWPI-Luc2 (a reporter plasmid encoding a firefly luciferase gene[13]) and 500 ng plasmids expressing parental S or its derivatives using TransIT-293 transfection reagent (Mirus, Cat# MIR2704) according to the manufacturer's protocol. Two days post transfection, the culture supernatants were harvested and filtrated. The amount of produced pseudovirus particles was quantified by the HiBiT assay using Nano Glo HiBiT lytic detection system (Promega, Cat# N3040) as previously described[13]. In this system, HiBiT peptide is produced with HIV-1 integrase and forms NanoLuc luciferase with LgBiT, which is supplemented with substrates. In each pseudovirus particle, the detected HiBiT value is correlated with the amount of the pseudovirus capsid protein, HIV-1 p24 protein.[13] Therefore, we calculated the amount of HIV-1 p24 capsid protein based on the HiBiT value measured, according to the previous paper. [13] To measure viral infectivity, the same amount of pseudovirus normalized with the HIV-1 p24 capsid protein was inoculated into HOS-ACE2/TMPRSS2 cells. At two days postinfection, the infected cells were lysed with a Bright-Glo luciferase assay system (Promega, Cat# E2620), and the luminescent signal produced by firefly luciferase reaction was measured using a GloMax explorer multimode microplate reader 3500 (Promega). The pseudoviruses were stored at –80°C until use.

**Neutralization assay**

Neutralization assays were performed previously described[5–8] with some modifications. First, the assays were mainly conducted by a semi-automated high-throughput method using Fluent780 (Tecan).[14] The SARS-CoV-2 spike pseudoviruses (counting ~100,000 relative light units) and serially diluted (40-fold to 29,160-fold dilution at the final concentration) heat-inactivated sera were manually prepared in a 2-ml 96-well plate (Greiner, Cat# 780271) and in 96-well microplates (ThermoFisher Scientific, Cat# 168136), respectively. The pseudoviruses were dispensed and mixed with the sera in 384-well plates (ThermoFisher Scientific, Cat# 164610) on Fluent780 (Tecan). Pseudoviruses without sera were included as controls. After incubation at 37°C for 1 hour, HOS-ACE2/TMPRSS2 cells (3,000 cells/30 μl) were added to the 20 μl mixture of pseudovirus and serum in the 384-well white plate on the device. Two days post infection, the infected cells were lysed with a Bright-Glo luciferase assay system (Promega, Cat# E2620) on Fluent780 (Tecan), and the luminescent signal was measured and processed using an Infinite200 and a Magellan (Tecan). The assay of each serum sample was performed in quadruplicate, and the 50% neutralization titer ($NT_{50}$) was calculated using Prism 9 (GraphPad Software).

**Data availability**

The GISAID datasets used in this study are available from the GISAID database (https://www.gisaid.org; EPI-SET-ID: EPI_SET_240418tb). The supplemental tables for the GISAID datasets are available in the GitHub repository (https://github.com/TheSatoLab/JN.1_short).

**Table S1. Human infection sera used in this study**

| SARS-CoV-2 infected | Donor ID | Sex | Age | Date of 1st vaccination (YYYY-MM-DD) | Date of 2nd vaccination (YYYY-MM-DD) | Date of 3rd vaccination (YYYY-MM-DD) | Date of 4th vaccination (YYYY-MM-DD) | Date of 5th vaccination (YYYY-MM-DD) | Date of 6th vaccination (YYYY-MM-DD) | Date of 7th vaccination (YYYY-MM-DD) | Date of test (YYYY-MM-DD) | Date of sampling (YYYY-MM-DD) | Prior infection? |
|---|---|---|---|---|---|---|---|---|---|---|---|---|---|
| XBB.1.5 | 37306 | Female | 53 | 2021-04-27 (P) | 2021-05-18 (P) | 2022-02-01 (P) | 2022-07-30 (M) | 2022-12-17 (P) | | | 2023-07-20 | 2023-08-08 | No |
| XBB.1.5 | 37097 | Female | 44 | NA (M) | 2021-08-16 (M) | 2022-05-13 (M) | | | | | 2023-07-13 | 2023-08-11 | No |
| XBB.1.5 | 37598 | Female | 43 | 2021-04-28 (P) | 2021-05-19 (P) | 2022-11-08 (P) | 2022-07-08 (M) | 2022-12-27 (P) | | | 2023-07-28 | 2023-08-11 | Yes |
| XBB.1.5 | 37072 | Female | 15 | 2021-09-25 (P) | 2021-10-18 (P) | 2022-05-02 (P) | | | | | 2023-07-11 | 2023-08-11 | No |
| XBB.1.5 | 37071 | Female | 48 | 2021-10-01 (P) | 2021-11-01 (P) | 2022-05-06 (P) | | | | | 2023-07-15 | 2023-08-11 | No |
| XBB.1.5 | 36845 | Male | 29 | 2021-09-01 (M) | 2021-09-29 (M) | 2022-05-27 (M) | | | | | 2023-06-26 | 2023-08-11 | No |
| XBB.1.5 | 37229 | Female | 74 | 2021-06-24 (P) | 2021-07-15 (P) | 2022-02-16 (M) | 2022-07-20 (M) | 2023-03-25 (M) | | | 2023-07-17 | 2023-08-01 | No |
| XBB.1.5 | 38084 | Male | 44 | 2021-09-13 (P) | 2021-10-05 (P) | 2022-07-29 (M) | | | | | 2023-08-11 | 2023-09-02 | No |
| XBB.1.5 | 37998 | Female | 65 | 2021-08-04 (P) | 2021-08-30 (P) | 2022-03-19 (M) | 2022-09-02 (P) | 2022-12-24 (P) | 2023-06-27 (PBA4/5) | | 2023-08-10 | 2023-09-02 | No |
| XBB.1.5 | 37798 | Female | 62 | 2021-03-17 (P) | 2021-04-09 (P) | 2021-12-23 (P) | 2022-07-28 (P) | 2023-06-17 (P) | | | 2023-08-03 | 2023-08-20 | No |
| XBB.1.5 | 38019 | Female | 18 | 2021-09-07 (P) | 2021-10-07 (P) | 2022-04-28 (P) | 2022-12-27 (P) | | | | 2023-08-10 | 2023-09-04 | No |
| XBB.1.5 | 38952 | Female | 54 | 2021-07-27 (M) | 2021-08-24 (M) | 2022-03-24 (P) | 2022-10-26 (P) | | | | 2023-08-23 | 2023-09-10 | No |
| XBB.1.5 | 38871 | Male | 46 | 2021-09-12 (P) | 2021-10-03 (P) | 2022-04-08 (M) | 2022-11-05 (M) | | | | 2023-08-22 | 2023-09-10 | No |
| XBB.1.5 | 38880 | Male | 51 | 2021-10-07 (P) | 2021-10-28 (P) | 2022-05-13 (M) | 2022-11-09 (PBA4/5) | | | | 2023-08-22 | 2023-09-16 | No |
| XBB.1.5 | 39018 | Female | 51 | 2021-07-29 (P) | 2021-08-23 (P) | 2022-03-19 (P) | | | | | 2023-08-25 | 2023-09-13 | Yes |
| XBB.1.5 | 39019 | Female | 22 | 2021-07-26 (M) | 2021-08-23 (M) | 2022-03-19 (P) | | | | | 2023-08-25 | 2023-09-13 | No |
| XBB.1.5 | 39296 | Male | 48 | 2021-07-15 (M) | 2021-08-23 (M) | 2022-03-24 (M) | 2022-12-25 (M) | | | | 2023-09-01 | 2023-09-23 | No |
| XBB.1.5 | 39502 | Female | 43 | 2021-07-31 (P) | 2021-08-23 (P) | 2022-03-15 (P) | 2022-11-01 (P) | | | | 2023-09-05 | 2023-09-26 | No |
| XBB.1.5 | 39321 | Female | 67 | 2021-06-14 (P) | 2021-07-05 (P) | 2022-02-10 (M) | 2022-07-26 (M) | 2023-01-06 (P) | | | 2023-09-01 | 2023-09-27 | No |
| XBB.1.5 | 37463 | Female | 21 | 2021-08 (M) | 2021-09 (M) | | | | | | 2023-07-24 | 2023-08-12 | No |
| EG.5 | P618 | Male | 54 | 2021-08-05 (P) | 2021-08-26 (P) | 2022-03-17 (P) | 2022-08-24 (P) | 2022-11-24 (P) | | | 2023-08-23 | 2023-09-07 | No |
| EG.5 | KK-230801 | Male | 56 | 2021-07-16 (P) | 2021-08-06 (P) | 2022-03-03 (M) | 2022-08-09 (P) | | | | 2023-07-28 | 2023-08-01 | No |
| EG.5 | 37301 | Female | 27 | 2021-08-01 (M) | 2022-03-06 (M) | | | | | | 2023-07-19 | 2023-08-12 | No |
| EG.5 | 37330 | Female | 42 | 2021-09-25 (P) | 2021-10-16 (P) | 2022-05-13 (P) | | | | | 2023-07-20 | 2023-08-13 | No |
| EG.5 | 38111 | Female | 50 | 2021-08-16 (P) | 2021-09-06 (P) | 2022-04-01 (P) | | | | | 2023-08-12 | 2023-09-02 | No |
| EG.5 | 38197 | Female | 49 | 2021-08-20 (P) | 2021-09-10 (P) | 2022-03-25 (M) | 2022-11-18 (P) | | | | 2023-08-13 | 2023-09-02 | No |
| EG.5 | 38217 | Male | 62 | 2021-07-04 (P) | 2021-07-25 (P) | 2022-02-26 (M) | 2022-08-06 (M) | 2022-11-20 (P) | | | 2023-08-13 | 2023-09-02 | No |
| EG.5 | 37999 | Male | 67 | 2021-06-07 (P) | 2021-07-01 (P) | 2022-02-08 (P) | 2022-07-12 (P) | 2022-12-24 (PBA4/5) | 2023-06-13 (MBA4/5) | | 2023-08-10 | 2023-09-02 | No |
| EG.5 | 38946 | Female | 65 | 2021-07-17 (P) | 2021-08-07 (P) | 2022-03-06 (M) | 2022-08-06 (M) | 2022-11-25 (P) | | | 2023-08-23 | 2023-09-10 | No |
| EG.5 | 39025 | Male | 63 | 2021-08-01 (P) | 2021-08-22 (P) | 2022-03-07 (M) | 2022-08-10 (M) | 2022-11-26 (PBA4/5) | 2023-06-06 (PBA4/5) | | 2023-08-26 | 2023-09-11 | No |
| EG.5 | 39288 | Male | 77 | 2021-06-09 (P) | 2021-07-01 (P) | 2022-02-05 (M) | 2022-07-12 (P) | 2022-11-18 (PBA4/5) | 2023-05-30 (PBA1) | | 2023-08-31 | 2023-09-20 | No |
| EG.5 | 39301 | Male | 41 | 2021-08-26 (M) | 2021-09-23 (M) | 2022-04-24 (M) | 2022-10-09 (MBA1) | | | | 2023-09-01 | 2023-09-23 | No |
| EG.5 | 39314 | Female | 56 | 2021-07-25 (P) | 2021-08-22 (P) | 2022-03-03 (M) | 2022-08-06 (M) | 2022-11-06 (M) | 2023-04-01 (NA) | | 2023-09-01 | 2023-09-21 | No |
| EG.5 | 39315 | Female | 60 | 2021-08-29 (P) | 2021-09-19 (P) | 2022-04-16 (M) | 2022-09-16 (P) | 2022-12-16 (PBA4/5) | | | 2023-08-31 | 2023-09-23 | No |
| EG.5 | 39316 | Male | 63 | 2021-07-31 (P) | 2021-08-21 (P) | 2022-04-23 (M) | 2022-09-30 (PBA1) | | | | 2023-08-31 | 2023-09-23 | No |
| EG.5 | 39330 | Male | 59 | 2021-08-24 (P) | 2021-09-14 (P) | 2022-03-26 (P) | 2022-10-23 (M) | | | | 2023-08-31 | 2023-09-23 | No |
| EG.5 | 39503 | Female | 41 | 2021-08-24 (P) | 2021-09-15 (P) | 2022-04-27 (P) | | | | | 2023-09-05 | 2023-09-25 | No |
| EG.5 | 39328 | Female | 59 | 2021-09-30 (P) | 2021-10-27 (P) | 2022-05-17 (P) | | | | | 2023-08-31 | 2023-09-27 | Yes |
| EG.5 | KK | Male | 43 | 2021-06-17 (P) | 2021-07-08 (P) | 2022-09-02 (P) | | | | | 2023-09-22 | 2023/10/03 | No |
| EG.5 | RK | Female | 41 | 2021-06-16 (P) | 2021-07-07 (P) | 2022-09-02 (P) | | | | | 2023-09-22 | 2023/10/03 | No |
| HK.3 | 42405 | Male | 56 | NA | 2021-08-27 (P) | 2022-03-14 (M) | 2022-08-22 (P) | 2022-12-01 (P) | 2023-07-26 (P) | | 2023-11-03 | 2024-02-17 | Yes |
| HK.3 | 42410 | Male | 66 | 2021-04-23 (P) | 2021-05-14 (P) | 2022-01-27 (P) | 2022-08-10 (P) | 2022-12-07 (M) | | | 2023-11-04 | 2024-02-13 | No |
| HK.3 | 42412 | Female | 75 | 2021-05-27 (P) | 2021-06-17 (P) | 2022-01-29 (P) | 2022-12-16 (P) | 2023-05-26 (P) | | | 2023-11-05 | 2024-02-18 | No |
| HK.3 | 42413 | Male | 10 | 2022-03-25 (P) | 2022-07-22 (P) | 2023-01-17 (P) | | | | | 2023-11-05 | 2024-02-18 | No |
| HK.3 | 42414 | Female | 41 | 2021-10-29 (P) | 2021-11-22 (P) | 2022-05-28 (P) | 2022-11-08 (P) | 2023-12-10 (P) | | | 2023-11-05 | 2024-02-18 | No |
| HK.3 | 42418 | Female | 25 | 2021-03-18 (P) | 2021-04-07 (P) | 2022-01-20 (P) | | | | | 2023-11-05 | 2024-02-16 | No |
| HK.3 | 42423 | Male | 49 | 2021-07-06 (M) | 2021-08-10 (M) | 2022-03-12 (M) | 2022-09-20 (M) | | | | 2023-11-05 | 2024-02-25 | No |
| HK.3 | 42429 | Male | 66 | 2021-07-05 (P) | 2021-07-26 (P) | 2022-03-04 (P) | | | | | 2023-11-06 | 2024-02-15 | NA |
| HK.3 | 39335 | Male | 50 | 2021-09-09 (P) | 2021-09-30 (P) | 2022-05-09 (P) | 2022-12-16 (P) | | | | 2023-08-31 | 2023-09-23 | NA |
| HK.3 | 39339-1 | Male | 53 | 2021-10-01 (P) | 2022-06-01 (P) | | | | | | 2023-09-02 | 2023-09-23 | NA |
| HK.3 | 3319 | Female | 55 | 2021-10-01 (P) | 2021-10-22 | 2022-07-10 | | | | | 2023-12-13 | 2024-01-17 | NA |
| HK.3 | 3363 | Male | 38 | NA | 2021-08 | | | | | | 2024-01-06 | 2024-01-27 | NA |
| HK.3 | 3373 | Male | 73 | 2022 | 2022 | | | | | | 2024-01-11 | 2024-02-01 | No |
| HK.3 | 3378 | Male | 75 | NA | 2021-06-11 | NA | | | | | 2024-01-14 | 2024-02-07 | NA |
| HK.3 | 3342 | Male | 76 | - | | | | | | | 2023-12-29 | 2024-01-19 | NA |
| HK.3 | 3353 | Male | 83 | - | | | | | | | 2024-01-02 | 2024-01-26 | NA |
| HK.3 | 3354 | Male | 84 | - | | | | | | | 2024-01-02 | 2024-01-19 | NA |
| HK.3 | 3391 | Male | 58 | - | | | | | | | 2024-01-20 | 2024-02-15 | NA |
| JN.1 | 3315 | Female | 75 | NA | NA | NA | NA | NA | NA | 2023-11-07 | 2023-12-11 | 2024-01-15 | No |
| JN.1 | 3323 | Female | 73 | 2021 | 2021 | 2021-05 | | | | | 2023-12-15 | 2023-12-28 | Yes |
| JN.1 | 3325 | Female | 73 | 2021-06 | 2021-07 | | | | | | 2023-12-16 | 2024-01-16 | NA |
| JN.1 | 3338 | Male | 31 | 2021-06 | 2021-07 | 2022-02-02 | | | | | 2023-12-26 | 2024-02-10 | Yes |
| JN.1 | 3355 | Male | 54 | NA | NA | NA | NA | NA | NA | NA | 2024-01-03 | 2024-02-15 | NA |
| JN.1 | 3316 | Female | 52 | - | | | | | | | 2023-12-11 | 2024-01-06 | NA |
| JN.1 | 3320 | Female | 80 | - | | | | | | | 2023-12-13 | 2024-01-12 | NA |
| JN.1 | 3329 | Male | 94 | - | | | | | | | 2023-12-18 | 2024-01-23 | NA |
| JN.1 | 3337 | Male | 72 | - | | | | | | | 2023-12-26 | 2024-01-19 | NA |
| JN.1 | 3362 | Male | 74 | - | | | | | | | 2024-01-06 | 2024-01-19 | NA |
| JN.1 | 3367 | Female | 70 | - | | | | | | | 2024-01-08 | 2024-01-26 | NA |
| JN.1 | 3400 | Female | 84 | - | | | | | | | 2024-01-24 | 2024-02-09 | NA |

NA, not applicable.
P, Pfizer-BioNTech; M, Moderna

**Table S2. Human XBB.1.5 vaccine sera used in this study**

| Donor ID | Sex | Age | Date of 1st vaccination (YYYY-MM-DD) | Date of 2nd vaccination (YYYY-MM-DD) | Date of 3rd vaccination (YYYY-MM-DD) | Date of 4th vaccination (YYYY-MM-DD) | Date of 5th vaccination (YYYY-MM-DD) | Date of 6th vaccination (YYYY-MM-DD) | Date of sampling (before vaccination) (YYYY-MM-DD) | Date of XBB.1.5 vaccination (YYYY-MM-DD) | Date of sampling (after vaccination) (YYYY-MM-DD) | Time interval between vaccination and the second sampling | Prior infection? | variant |
|---|---|---|---|---|---|---|---|---|---|---|---|---|---|---|
| 5165 | Female | 89 | 2021-05-29 (P) | 2021-06-21 (P) | 2022-02-16 (P) | 2022-07-17 (P) | 2022-11-27 (PBA4/5) | 2023-05-20 (PBA4/5) | 2023-09-29 | 2023-09-29 (XBB1.5) | 2023-10-26 | 27 | No | |
| 5166 | Male | 77 | 2021-06-05 (P) | 2021-07-04 (P) | 2022-03-05 (P) | 2022-08-06 (P) | 2022-11-20 (PBA4/5) | 2023-05-20 (PBA4/5) | 2023-09-29 | 2023-09-29 (XBB1.5) | 2023-10-21 | 22 | No | |
| 6783 | Male | 57 | 2021-06-23 (M) | 2021-07-21 (M) | 2022-02-11 (M) | 2022-10-15 (MBA1) | | | 2023-10-03 | 2023-10-03 (XBB1.5) | 2023-10-23 | 20 | No | |
| 2477 | Male | 81 | 2021-07-10 (P) | 2021-07-31 (P) | 2022-03-06 (P) | 2022-08-06 (P) | 2022-11-07 (PBA4/5) | 2023-05-09 (PBA4/5) | 2023-09-29 | 2023-09-29 (XBB1.5) | 2023-10-28 | 29 | No | |
| 6858 | Female | 62 | 2021-07-18 (P) | 2021-08-11 (P) | 2022-02-27 (M) | 2022-07-30 (P) | 2022-11-20 (PBA4/5) | | 2023-10-07 | 2023-10-07 (XBB1.5) | 2023-10-28 | 21 | No | |
| 192 | Female | 64 | 2021-04-22 (P) | 2021-05-13 (P) | 2022-01-15 (P) | 2022-07-16 (P) | 2022-11-24 (PBA4/5) | 2023-05-25 (PBA4/5) | 2023-10-02 | 2023-10-26 (PXBB1.5) | 2023-11-20 | 25 | No | |
| 1700 | Female | 53 | 2021-04-21 (P) | 2021-05-12 (P) | 2022-01-15 (P) | 2022-07-13 (P) | 2022-11-30 (PBA4/5) | 2023-06-23 (MBA4/5) | 2023-09-29 | 2023-10-18 (PXBB1.5) | 2023-11-13 | 26 | No | |
| 5555 | Female | 51 | 2021-07-14 (P) | 2021-08-14 (P) | 2022-02-22 (P) | 2022-07-23 (P) | 2022-12-03 (PBA4/5) | 2023-05-11 (PBA4/5) | 2023-09-30 | 2023-10-21 (PXBB1.5) | 2023-11-15 | 25 | No | |
| 5986 | Male | 62 | 2021-07-24 (P) | 2021-08-14 (P) | 2022-03-12 (M) | 2022-08-27 (M) | 2022-12-24 (PBA4/5) | 2023-06-03 (PBA4/5) | 2023-09-25 | 2023-10-21 (PXBB1.5) | 2023-11-13 | 23 | No | |
| KS | Male | 41 | 2021-06-17 (P) | 2021-07-07 (P) | 2022-03-28 (M) | 2022-10-27 (MBA.5) | | | 2023-09-19 | 2023-09-20 (XBB1.5) | 2023-10-14 | 24 | Yes (2023-06-29) | XBB.1.9 |
| KY | Female | 53 | 2021-08-18 (P) | 2021-09-08 (P) | 2022-04-13 (P) | 2022-10-21 (P) | | | 2023-09-25 | 2023-09-27 (XBB1.5) | 2023-10-20 | 23 | Yes (2023-07-24) | XBB.1.16 |
| KK | Male | 56 | 2021-07-16 (P) | 2021-08-06 (P) | 2022-03-03 (M) | 2022-08-09 (P) | | | 2023-09-25 | 2023-09-29 (XBB1.5) | 2023-10-24 | 25 | Yes (2023-07-17) | EG.5 |
| 2345 | Female | 52 | 2021-07-11 (P) | 2021-08-01 (P) | 2022-03-10 (P) | 2022-10-22 (PBA1) | | | 2023-10-06 | 2023-10-06 (XBB1.5) | 2023-10-27 | 21 | Yes (2023-07) | NA |
| 80 | Male | 61 | 2021-05-10 (P) | 2021-05-31 (P) | 2022-01-24 (M) | 2022-07-22 (BA4/5) | 2022-11-12 (BA4/5) | 2023-06-03 (BA4/5) | 2023-09-29 | 2023-09-30 (XBB1.5) | 2023-10-24 | 24 | Yes (2023-07) | NA |
| 90 | Male | 47 | 2021-05-11 (P) | 2021-06-02 (P) | 2022-01-25 (M) | 2022-08-02 (BA4/5) | 2022-11-09 (BA4/5) | | 2023-10-10 | 2023-10-11 (XBB1.5) | 2023-11-01 | 21 | Yes (2023-07) | NA |
| 100 | Male | 32 | 2021-05-18 (P) | 2021-06-08 (P) | 2022-01-31 (M) | 2022-07-17 (MBA4/5) | 2022-11-15 (PBA4/5) | 2023-05-27 (MBA4/5) | 2023-10-11 | 2023-10-14 (XBB1.5) | 2023-11-06 | 23 | Yes (2023-06) | NA |
| 286691 | Male | 49 | 2021-06-01(P) | 2021-06-22 (P) | 2022-03-21 (M) | | | | 2023-10-19 | 2023-10-19 (PXBB1.5) | 2023-11-16 | 28 | Yes (2023-08-22) | XBB.1.9 |
| 2439306 | Male | 43 | 2021-03-08 (P) | 2021-03-29 (P) | 2021-12-20 (P) | 2022-08-26 (M) | 2022-11-30 (M) | | 2023-10-23 | 2023-10-23 (PXBB1.5) | 2023-11-20 | 28 | Yes (2023-09-08) | EG.5 |
| 4177 | Female | 49 | 2021-04-24 (P) | 2021-05-15 (P) | 2022-01-22 (P) | 2022-07-09 (P) | 2022-12-03 (PBA4/5) | 2023-06-13 (MBA4/5) | 2023-09-29 | 2023-10-14 (PXBB1.5) | 2023-11-10 | 27 | Yes (2023-08-29) | NA |
| 38579 | Male | 48 | 2022-05-28 (NA) | | | | | | 2023-10-31 | 2023-10-31 (XBB1.5) | 2023-11-26 | 26 | Yes (2023-08-16) | XBB.1.16 |
| 36708 | Male | 55 | 2021-08-07 (P) | 2021-08-27 (P) | 2022-04-14 (P) | | | | 2023-10-21 | 2023-11-11 (XBB1.5) | 2023-12-09 | 28 | Yes (2023-05-31) | XBB.1.5 |
| 38061 | Female | 55 | 2021-08-17 (P) | 2021-09-18 (P) | 2022-04-02 (M) | 2022-10-14 (PBA1) | | | 2023-11-28 | 2023-11-28 (XBB1.5) | 2023-12-19 | 21 | Yes (2023-08-10) | XBB.1.5 |

NA, not applicable.

P, Pfizer-BioNTech; M, Moderna

**Table S3. Estimated relative Re and epidemic dynamics modeling parameters of the representative SARS-CoV-2 Omicron sublineages spreading in USA, United Kingdom, and Canada from Nov 1, 2023 to April 1, 2023**

| | | Relative $R_e$ (posterior values) | | | | Bulk effective | Tail effective |
|---|---|---|---|---|---|---|---|
| PANGO lineage | Country | Mean | $2.5^{th}$ percentile | $97.5^{th}$ percentile | $\hat{R}$ | sample size | sample size |
| KP.2 | USA | 1.224 | 1.193 | 1.256 | 1.000 | 21114.1 | 11075.1 |
| KP.1.1 | USA | 1.191 | 1.145 | 1.243 | 1.000 | 20045.9 | 11192.3 |
| JN.1.16.1 | USA | 1.179 | 1.143 | 1.218 | 1.000 | 20799.9 | 11970.9 |
| JN.1+F456L | USA | 1.088 | 1.079 | 1.098 | 1.001 | 22614.1 | 12052.3 |
| XDP.1 | USA | 1.080 | 1.058 | 1.102 | 1.001 | 22413.1 | 11623.1 |
| JN.1+R346T | USA | 1.070 | 1.065 | 1.074 | 1.000 | 17870.7 | 12073.4 |
| XDK | USA | 1.063 | 1.040 | 1.086 | 1.000 | 21764.0 | 11994.5 |
| XDQ | USA | 1.050 | 1.030 | 1.071 | 1.000 | 22467.8 | 11694.9 |
| XDP | USA | 1.019 | 1.010 | 1.028 | 1.001 | 22688.3 | 11904.6 |
| JN.2.5 | USA | 0.997 | 0.972 | 1.023 | 1.000 | 21902.3 | 12207.9 |
| XDD | USA | 0.988 | 0.978 | 0.999 | 1.000 | 23476.2 | 12692.4 |
| BA.2 | USA | 0.951 | 0.938 | 0.964 | 1.000 | 22196.8 | 11728.2 |
| GE.1.2 | USA | 0.950 | 0.927 | 0.972 | 1.000 | 25385.4 | 11655.7 |
| JN.4 | USA | 0.937 | 0.913 | 0.960 | 1.000 | 24757.1 | 12044.8 |
| JN.6 | USA | 0.920 | 0.893 | 0.947 | 1.000 | 23495.7 | 12668.9 |
| BA.2.86.1 | USA | 0.918 | 0.911 | 0.925 | 1.000 | 21009.5 | 10926.1 |
| BA.2.86 | USA | 0.913 | 0.891 | 0.935 | 1.000 | 23459.5 | 12086.5 |
| GW.5 | USA | 0.913 | 0.888 | 0.938 | 1.000 | 23624.5 | 12511.2 |
| JN.2 | USA | 0.908 | 0.897 | 0.918 | 1.000 | 22214.4 | 11979.3 |
| JN.11 | USA | 0.905 | 0.878 | 0.932 | 1.000 | 22242.7 | 13286.8 |
| JN.3 | USA | 0.900 | 0.892 | 0.909 | 1.000 | 20357.3 | 12669.4 |
| XBB.1.41.1 | USA | 0.898 | 0.885 | 0.910 | 1.000 | 22994.9 | 11600.0 |
| JD.1.1.1 | USA | 0.896 | 0.888 | 0.903 | 1.000 | 21085.0 | 12089.5 |
| JG.3 | USA | 0.891 | 0.888 | 0.895 | 1.000 | 14447.2 | 11928.0 |
| FL.15.1.1 | USA | 0.889 | 0.875 | 0.903 | 1.000 | 25319.3 | 12229.3 |
| JD.1.1.3 | USA | 0.886 | 0.868 | 0.903 | 1.000 | 21966.8 | 11953.2 |
| XBB.1.42.1 | USA | 0.883 | 0.850 | 0.916 | 1.000 | 21781.8 | 12131.9 |
| HK.3.2 | USA | 0.882 | 0.874 | 0.889 | 1.000 | 20188.5 | 12596.6 |
| JE.1.1 | USA | 0.879 | 0.864 | 0.895 | 1.001 | 21934.4 | 12609.8 |
| JQ.1 | USA | 0.878 | 0.852 | 0.904 | 1.000 | 23216.0 | 11775.8 |
| JE.1 | USA | 0.878 | 0.854 | 0.901 | 1.000 | 21333.0 | 11678.6 |
| XBB.1.16.17 | USA | 0.877 | 0.868 | 0.886 | 1.000 | 21132.6 | 12114.6 |
| FL.36 | USA | 0.877 | 0.845 | 0.909 | 1.000 | 23896.1 | 12154.1 |
| HK.3.1 | USA | 0.873 | 0.859 | 0.887 | 1.000 | 21163.1 | 12286.3 |
| JD.1.1 | USA | 0.872 | 0.868 | 0.876 | 1.000 | 15803.9 | 11651.8 |
| HK.11 | USA | 0.869 | 0.846 | 0.890 | 1.000 | 21925.2 | 12197.8 |
| FY.5 | USA | 0.868 | 0.855 | 0.880 | 1.001 | 20697.2 | 12499.9 |

| | | | | | | | |
|---|---|---|---|---|---|---|---|
| FL.1.5.2 | USA | 0.868 | 0.857 | 0.878 | 1.000 | 19865.4 | 12262.6 |
| JJ.1 | USA | 0.866 | 0.836 | 0.896 | 1.000 | 22242.7 | 12208.6 |
| FL.13 | USA | 0.866 | 0.831 | 0.899 | 1.001 | 23424.4 | 12744.8 |
| FL.20.2 | USA | 0.864 | 0.840 | 0.887 | 1.001 | 19369.4 | 12374.3 |
| HV.1 | USA | 0.860 | 0.858 | 0.862 | 1.001 | 7465.2 | 9761.9 |
| HK.1 | USA | 0.860 | 0.844 | 0.875 | 1.000 | 19796.5 | 11890.5 |
| HK.8 | USA | 0.858 | 0.840 | 0.875 | 1.000 | 21846.3 | 13010.9 |
| GK.1 | USA | 0.857 | 0.846 | 0.868 | 1.000 | 20819.2 | 12441.4 |
| GS.4 | USA | 0.856 | 0.830 | 0.882 | 1.000 | 21811.2 | 13148.6 |
| XBB.2.3.3 | USA | 0.854 | 0.815 | 0.891 | 1.001 | 22780.3 | 12285.4 |
| HK.3 | USA | 0.853 | 0.849 | 0.857 | 1.000 | 14533.6 | 11594.7 |
| GK.1.3 | USA | 0.853 | 0.839 | 0.867 | 1.000 | 21261.9 | 12420.8 |
| EG.5.1.8 | USA | 0.853 | 0.845 | 0.861 | 1.001 | 20945.6 | 11775.6 |
| XBB.1.5 | USA | 0.853 | 0.833 | 0.871 | 1.000 | 21234.8 | 11390.0 |
| HK.2 | USA | 0.852 | 0.828 | 0.875 | 1.000 | 22539.0 | 12483.2 |
| GS.4.1 | USA | 0.851 | 0.840 | 0.862 | 1.000 | 19979.5 | 12730.2 |
| GK.2.1 | USA | 0.851 | 0.821 | 0.881 | 1.000 | 22092.3 | 12142.4 |
| GA.4.1 | USA | 0.851 | 0.813 | 0.887 | 1.000 | 23880.0 | 11676.8 |
| JD.1 | USA | 0.851 | 0.815 | 0.884 | 1.000 | 21173.4 | 12632.5 |
| JF.1 | USA | 0.847 | 0.841 | 0.853 | 1.000 | 19999.6 | 11492.9 |
| GK.1.4 | USA | 0.845 | 0.807 | 0.881 | 1.001 | 21974.6 | 12214.0 |
| GK.1.1 | USA | 0.845 | 0.837 | 0.853 | 1.000 | 19039.7 | 11909.4 |
| XBB.1.16.15 | USA | 0.844 | 0.836 | 0.853 | 1.000 | 20695.6 | 12814.0 |
| XBB.1 | USA | 0.844 | 0.811 | 0.876 | 1.000 | 20554.9 | 12425.3 |
| XBB.2.3 | USA | 0.842 | 0.822 | 0.862 | 1.000 | 24847.4 | 12062.2 |
| XBB.1.41.2 | USA | 0.840 | 0.817 | 0.861 | 1.000 | 23444.3 | 13067.4 |
| EG.5.1.6 | USA | 0.839 | 0.832 | 0.845 | 1.000 | 18456.3 | 12588.9 |
| HF.1.2 | USA | 0.838 | 0.800 | 0.874 | 1.000 | 22482.0 | 12142.7 |
| JC.1 | USA | 0.836 | 0.798 | 0.873 | 1.000 | 20050.9 | 11934.1 |
| GN.1 | USA | 0.836 | 0.803 | 0.868 | 1.000 | 21133.3 | 13137.9 |
| EG.5.1.1 | USA | 0.836 | 0.831 | 0.841 | 1.000 | 16127.6 | 11768.6 |
| FU.2 | USA | 0.834 | 0.810 | 0.857 | 1.000 | 21647.6 | 12400.3 |
| HF.1.1 | USA | 0.833 | 0.819 | 0.847 | 1.000 | 23045.8 | 12467.7 |
| HN.1 | USA | 0.833 | 0.815 | 0.851 | 1.000 | 23854.2 | 12174.3 |
| DV.7.1 | USA | 0.832 | 0.810 | 0.854 | 1.000 | 24359.9 | 12262.7 |
| GK.2 | USA | 0.831 | 0.808 | 0.854 | 1.000 | 22698.6 | 12899.3 |
| EG.5.1 | USA | 0.831 | 0.825 | 0.837 | 1.000 | 16769.8 | 11651.9 |
| HK.6 | USA | 0.831 | 0.811 | 0.849 | 1.000 | 25973.9 | 12124.2 |
| HK.3.3 | USA | 0.830 | 0.801 | 0.857 | 1.000 | 23139.5 | 12522.7 |
| XBB.1.16.11 | USA | 0.829 | 0.822 | 0.836 | 1.000 | 18399.5 | 12599.0 |
| GJ.1.2.2 | USA | 0.828 | 0.807 | 0.848 | 1.000 | 22892.1 | 12509.0 |

| | | | | | | | |
|---|---|---|---|---|---|---|---|
| GE.1 | USA | 0.826 | 0.811 | 0.840 | 1.001 | 19467.0 | 12612.9 |
| XBB.1.16.6 | USA | 0.825 | 0.820 | 0.831 | 1.000 | 17875.7 | 12207.9 |
| XBB.1.42.2 | USA | 0.825 | 0.802 | 0.847 | 1.000 | 21431.2 | 12568.8 |
| EG.10.1 | USA | 0.824 | 0.797 | 0.851 | 1.000 | 20566.6 | 12489.4 |
| FL.1.5.1 | USA | 0.823 | 0.819 | 0.828 | 1.000 | 15101.4 | 11632.8 |
| XCH.1 | USA | 0.823 | 0.799 | 0.847 | 1.001 | 24047.0 | 12624.0 |
| XBB.1.9.1 | USA | 0.823 | 0.793 | 0.853 | 1.001 | 21661.6 | 12368.0 |
| EG.5 | USA | 0.823 | 0.799 | 0.846 | 1.000 | 23483.5 | 12709.7 |
| FL.24 | USA | 0.823 | 0.780 | 0.863 | 1.000 | 23749.6 | 11818.0 |
| XBB.1.9 | USA | 0.822 | 0.799 | 0.843 | 1.000 | 23911.4 | 13209.0 |
| HF.1 | USA | 0.819 | 0.806 | 0.831 | 1.000 | 21232.6 | 12947.9 |
| GK.3.1 | USA | 0.817 | 0.786 | 0.846 | 1.001 | 23397.4 | 12083.2 |
| EG.5.1.3 | USA | 0.815 | 0.805 | 0.825 | 1.000 | 21766.9 | 12430.8 |
| HZ.1 | USA | 0.815 | 0.788 | 0.841 | 1.000 | 22002.1 | 11481.4 |
| XBB.1.16 | USA | 0.815 | 0.798 | 0.831 | 1.001 | 22770.7 | 12736.4 |
| XBB.1.5.15 | USA | 0.812 | 0.774 | 0.848 | 1.000 | 21076.6 | 12007.8 |
| EG.5.1.4 | USA | 0.810 | 0.800 | 0.820 | 1.000 | 21612.0 | 12284.4 |
| XBB.1.16.9 | USA | 0.807 | 0.781 | 0.832 | 1.000 | 26892.6 | 12174.9 |
| EG.6.1 | USA | 0.801 | 0.776 | 0.824 | 1.000 | 24454.1 | 12388.8 |
| XCH | USA | 0.800 | 0.772 | 0.826 | 1.000 | 22798.5 | 13115.7 |
| EG.6.1.1 | USA | 0.798 | 0.760 | 0.835 | 1.000 | 22205.7 | 13036.7 |
| XBB.1.5.70 | USA | 0.796 | 0.761 | 0.828 | 1.001 | 23270.2 | 13409.1 |
| XBB.1.9.2 | USA | 0.795 | 0.771 | 0.818 | 1.000 | 24098.9 | 11187.4 |
| XBB.1.16.23 | USA | 0.794 | 0.768 | 0.819 | 1.000 | 25558.7 | 12230.5 |
| GJ.1.2 | USA | 0.790 | 0.769 | 0.810 | 1.000 | 19870.2 | 12893.1 |
| FU.1 | USA | 0.777 | 0.735 | 0.814 | 1.000 | 21411.8 | 12680.9 |
| XBB.1.16.1 | USA | 0.765 | 0.713 | 0.814 | 1.000 | 20919.0 | 14031.8 |
| FL.20 | USA | 0.763 | 0.722 | 0.802 | 1.000 | 21747.2 | 12604.9 |
| KP.2 | United Kingdom | 1.320 | 1.237 | 1.423 | 1.001 | 30172.2 | 11665.8 |
| JN.1.16.1 | United Kingdom | 1.246 | 1.181 | 1.327 | 1.000 | 29303.8 | 11259.9 |
| JN.1+F456L | United Kingdom | 1.080 | 1.066 | 1.095 | 1.001 | 35263.4 | 11720.7 |
| XDK | United Kingdom | 1.077 | 1.060 | 1.096 | 1.000 | 36202.3 | 11652.9 |
| JN.1+R346T | United Kingdom | 1.065 | 1.057 | 1.074 | 1.000 | 28730.4 | 11005.6 |
| XDS | United Kingdom | 1.056 | 1.031 | 1.082 | 1.000 | 31963.4 | 11470.4 |
| XDT | United Kingdom | 1.044 | 1.012 | 1.077 | 1.001 | 31209.8 | 11989.8 |
| XDN | United Kingdom | 1.000 | 0.988 | 1.013 | 1.001 | 33186.0 | 10792.7 |
| JN.10 | United Kingdom | 0.979 | 0.954 | 1.004 | 1.000 | 33356.2 | 11074.5 |
| XDD | United Kingdom | 0.969 | 0.949 | 0.988 | 1.000 | 34457.2 | 10779.1 |
| JN.4 | United Kingdom | 0.961 | 0.943 | 0.980 | 1.000 | 27604.8 | 11640.2 |
| JN.6 | United Kingdom | 0.925 | 0.915 | 0.935 | 1.000 | 25516.6 | 12490.7 |
| JN.2 | United Kingdom | 0.922 | 0.914 | 0.930 | 1.000 | 24957.3 | 11736.3 |

| | | | | | | | |
|---|---|---|---|---|---|---|---|
| JN.9 | United Kingdom | 0.917 | 0.888 | 0.945 | 1.000 | 33105.8 | 11610.8 |
| JD.1.1.1 | United Kingdom | 0.909 | 0.884 | 0.933 | 1.000 | 34450.2 | 11522.5 |
| BA.2.86.3 | United Kingdom | 0.907 | 0.884 | 0.930 | 1.000 | 29146.2 | 10584.2 |
| BA.2.86.1 | United Kingdom | 0.904 | 0.897 | 0.910 | 1.000 | 19497.9 | 12400.1 |
| JN.3 | United Kingdom | 0.903 | 0.892 | 0.913 | 1.001 | 26262.6 | 11386.2 |
| XBB.1.16.17 | United Kingdom | 0.902 | 0.888 | 0.915 | 1.001 | 27274.1 | 11528.3 |
| BA.2.86.5 | United Kingdom | 0.900 | 0.863 | 0.936 | 1.000 | 32200.3 | 11673.2 |
| JN.5 | United Kingdom | 0.892 | 0.865 | 0.918 | 1.000 | 31123.1 | 11448.4 |
| HK.3.1 | United Kingdom | 0.890 | 0.862 | 0.917 | 1.000 | 31067.5 | 11919.3 |
| XBB.1.41.1 | United Kingdom | 0.877 | 0.857 | 0.897 | 1.000 | 28428.7 | 12417.2 |
| GW.5 | United Kingdom | 0.876 | 0.845 | 0.907 | 1.000 | 30977.9 | 11926.3 |
| JD.1.1 | United Kingdom | 0.876 | 0.866 | 0.885 | 1.000 | 21732.0 | 12018.3 |
| EG.5.1 | United Kingdom | 0.871 | 0.856 | 0.886 | 1.000 | 25151.4 | 11525.2 |
| GS.4.1 | United Kingdom | 0.870 | 0.856 | 0.884 | 1.001 | 28475.3 | 11634.8 |
| HK.3.2 | United Kingdom | 0.867 | 0.847 | 0.886 | 1.001 | 32635.9 | 11959.0 |
| JG.3 | United Kingdom | 0.864 | 0.855 | 0.873 | 1.000 | 20067.0 | 12521.1 |
| HV.1 | United Kingdom | 0.863 | 0.853 | 0.872 | 1.001 | 21985.7 | 12823.4 |
| FL.15.1.1 | United Kingdom | 0.862 | 0.823 | 0.899 | 1.000 | 31894.4 | 11815.2 |
| FL.20.1 | United Kingdom | 0.856 | 0.820 | 0.892 | 1.000 | 30848.5 | 10689.3 |
| FY.5 | United Kingdom | 0.852 | 0.815 | 0.887 | 1.000 | 29934.8 | 12371.1 |
| HK.3 | United Kingdom | 0.849 | 0.839 | 0.859 | 1.000 | 20399.5 | 11230.7 |
| EG.5.1.8 | United Kingdom | 0.848 | 0.827 | 0.868 | 1.000 | 27432.6 | 11331.6 |
| BA.2.86.2 | United Kingdom | 0.848 | 0.821 | 0.874 | 1.001 | 32964.2 | 11641.7 |
| GA.4.1 | United Kingdom | 0.832 | 0.790 | 0.872 | 1.000 | 28725.0 | 11726.6 |
| HF.1 | United Kingdom | 0.823 | 0.790 | 0.855 | 1.000 | 28400.8 | 11343.3 |
| FL.1.5.1 | United Kingdom | 0.821 | 0.803 | 0.839 | 1.000 | 28019.3 | 11743.9 |
| EG.5.1.3 | United Kingdom | 0.821 | 0.800 | 0.842 | 1.000 | 28147.8 | 12409.8 |
| JF.1 | United Kingdom | 0.820 | 0.800 | 0.839 | 1.000 | 27874.7 | 12213.0 |
| FL.1.5.2 | United Kingdom | 0.818 | 0.792 | 0.843 | 1.000 | 28461.7 | 11954.3 |
| EG.5.1.6 | United Kingdom | 0.817 | 0.795 | 0.838 | 1.000 | 27678.8 | 11487.2 |
| XBB.1.16.11 | United Kingdom | 0.815 | 0.795 | 0.835 | 1.000 | 26541.6 | 12136.0 |
| GK.1.1 | United Kingdom | 0.812 | 0.777 | 0.846 | 1.000 | 27796.2 | 11141.1 |
| EG.5.1.1 | United Kingdom | 0.809 | 0.795 | 0.822 | 1.000 | 22296.0 | 12392.0 |
| XBB.1.16 | United Kingdom | 0.807 | 0.772 | 0.840 | 1.000 | 30711.0 | 12244.9 |
| EG.5.1.4 | United Kingdom | 0.802 | 0.766 | 0.837 | 1.000 | 28375.9 | 11500.2 |
| XBB.1.16.6 | United Kingdom | 0.796 | 0.774 | 0.817 | 1.000 | 27018.5 | 11390.9 |
| XBB.1.16.15 | United Kingdom | 0.795 | 0.774 | 0.815 | 1.001 | 25025.5 | 11672.0 |
| HK.1 | United Kingdom | 0.791 | 0.746 | 0.834 | 1.000 | 30208.9 | 12388.3 |
| HK.6 | United Kingdom | 0.791 | 0.765 | 0.816 | 1.000 | 27266.0 | 13419.4 |
| HK.11 | United Kingdom | 0.782 | 0.731 | 0.829 | 1.000 | 31087.5 | 11841.0 |
| GE.1 | United Kingdom | 0.781 | 0.743 | 0.817 | 1.000 | 28678.5 | 11695.4 |

| | | | | | | | |
|---|---|---|---|---|---|---|---|
| EG.10.1 | United Kingdom | 0.774 | 0.741 | 0.805 | 1.000 | 29765.6 | 12293.5 |
| GK.2.3 | United Kingdom | 0.743 | 0.684 | 0.797 | 1.000 | 28541.0 | 13060.2 |
| KP.2 | Canada | 1.255 | 1.190 | 1.328 | 1.001 | 22517.1 | 11575.7 |
| JN.1+F456L | Canada | 1.171 | 1.155 | 1.188 | 1.000 | 22520.5 | 11160.3 |
| JN.1+R346T | Canada | 1.094 | 1.081 | 1.107 | 1.001 | 21206.5 | 11186.3 |
| XDK | Canada | 1.067 | 1.032 | 1.105 | 1.001 | 20697.6 | 10996.7 |
| XDP | Canada | 1.053 | 1.033 | 1.073 | 1.000 | 19844.3 | 11699.2 |
| XDD | Canada | 0.988 | 0.967 | 1.009 | 1.000 | 18303.4 | 11618.1 |
| JN.2.5 | Canada | 0.986 | 0.980 | 0.991 | 1.000 | 17497.9 | 11715.8 |
| JE.1.1 | Canada | 0.912 | 0.890 | 0.934 | 1.000 | 18046.1 | 11472.9 |
| JG.3 | Canada | 0.904 | 0.900 | 0.908 | 1.000 | 10602.3 | 11350.2 |
| BA.2.86.1 | Canada | 0.900 | 0.890 | 0.910 | 1.000 | 19330.3 | 11579.5 |
| JD.1.1.3 | Canada | 0.898 | 0.874 | 0.922 | 1.000 | 20131.8 | 11684.9 |
| GK.2.1 | Canada | 0.892 | 0.858 | 0.924 | 1.002 | 20540.2 | 12512.9 |
| JN.2 | Canada | 0.889 | 0.881 | 0.898 | 1.001 | 17536.9 | 12390.8 |
| XBB.1.5 | Canada | 0.889 | 0.860 | 0.917 | 1.000 | 19730.4 | 11928.1 |
| EG.5.1.8 | Canada | 0.875 | 0.861 | 0.888 | 1.000 | 19918.0 | 12200.9 |
| JD.1.1.1 | Canada | 0.875 | 0.866 | 0.883 | 1.000 | 15907.9 | 12120.4 |
| HR.1 | Canada | 0.872 | 0.843 | 0.901 | 1.000 | 21585.4 | 11716.4 |
| FL.1.5.2 | Canada | 0.871 | 0.861 | 0.881 | 1.001 | 16110.4 | 11921.0 |
| FY.5 | Canada | 0.865 | 0.844 | 0.884 | 1.000 | 20034.1 | 12900.2 |
| XBB.1.16.17 | Canada | 0.862 | 0.840 | 0.883 | 1.000 | 23978.2 | 12140.1 |
| HK.3.1 | Canada | 0.861 | 0.833 | 0.889 | 1.001 | 20149.0 | 10240.4 |
| JD.1.1 | Canada | 0.859 | 0.853 | 0.866 | 1.000 | 13890.4 | 12386.0 |
| HC.2 | Canada | 0.856 | 0.818 | 0.894 | 1.000 | 20161.5 | 11899.9 |
| EG.5.1.4 | Canada | 0.852 | 0.843 | 0.861 | 1.000 | 16925.9 | 13004.7 |
| HK.3 | Canada | 0.852 | 0.846 | 0.857 | 1.000 | 12068.8 | 11086.3 |
| HV.1 | Canada | 0.849 | 0.846 | 0.852 | 1.001 | 6497.4 | 10183.9 |
| JN.3 | Canada | 0.848 | 0.821 | 0.875 | 1.000 | 21882.6 | 11613.1 |
| EG.5.1.6 | Canada | 0.846 | 0.835 | 0.857 | 1.000 | 18882.0 | 12265.0 |
| HK.3.2 | Canada | 0.846 | 0.835 | 0.856 | 1.001 | 17607.8 | 12716.2 |
| GK.1 | Canada | 0.844 | 0.824 | 0.863 | 1.000 | 18406.1 | 12241.2 |
| FL.15.1.1 | Canada | 0.844 | 0.819 | 0.867 | 1.000 | 21206.3 | 12504.1 |
| XCH | Canada | 0.834 | 0.807 | 0.860 | 1.000 | 21288.1 | 11830.2 |
| DV.7.1 | Canada | 0.833 | 0.821 | 0.846 | 1.001 | 18651.6 | 12569.8 |
| EG.5.1.1 | Canada | 0.832 | 0.825 | 0.839 | 1.000 | 13884.1 | 12407.4 |
| DV.7.1.2 | Canada | 0.830 | 0.787 | 0.870 | 1.000 | 20515.5 | 12151.1 |
| XBB.1.16.11 | Canada | 0.829 | 0.802 | 0.855 | 1.000 | 21947.4 | 12473.3 |
| GK.1.1 | Canada | 0.828 | 0.812 | 0.843 | 1.000 | 20267.2 | 12777.0 |
| HK.6 | Canada | 0.827 | 0.802 | 0.851 | 1.000 | 21035.4 | 11995.8 |
| JF.1 | Canada | 0.824 | 0.811 | 0.836 | 1.000 | 18139.4 | 12352.2 |

| | | | | | | | |
|---|---|---|---|---|---|---|---|
| GA.4.1 | Canada | 0.816 | 0.794 | 0.839 | 1.000 | 19518.6 | 12066.3 |
| XBB.1.16.15 | Canada | 0.816 | 0.787 | 0.844 | 1.000 | 20176.2 | 13095.5 |
| EG.5.1 | Canada | 0.808 | 0.798 | 0.818 | 1.000 | 16799.1 | 12994.4 |
| GK.2 | Canada | 0.805 | 0.780 | 0.830 | 1.001 | 21684.6 | 12171.0 |
| XCH.1 | Canada | 0.804 | 0.762 | 0.844 | 1.000 | 18092.4 | 12196.0 |
| XBB.1.16.6 | Canada | 0.802 | 0.790 | 0.814 | 1.000 | 18305.8 | 12181.0 |
| FL.1.5.1 | Canada | 0.800 | 0.789 | 0.811 | 1.000 | 16851.3 | 11931.3 |
| GK.3.1 | Canada | 0.788 | 0.751 | 0.825 | 1.000 | 20882.7 | 12166.4 |
| GS.4.1 | Canada | 0.780 | 0.749 | 0.808 | 1.000 | 21371.0 | 13261.4 |
| XBB.1.16 | Canada | 0.778 | 0.748 | 0.807 | 1.000 | 22981.6 | 11553.8 |
| EG.5.1.3 | Canada | 0.776 | 0.759 | 0.793 | 1.000 | 19387.7 | 12236.3 |
| FL.15 | Canada | 0.770 | 0.724 | 0.813 | 1.000 | 20982.8 | 11748.1 |
| EG.5.2.1 | Canada | 0.763 | 0.711 | 0.811 | 1.000 | 21420.6 | 12883.7 |
| GJ.1.2 | Canada | 0.739 | 0.690 | 0.783 | 1.000 | 20201.9 | 11494.5 |
| EG.6.1 | Canada | 0.734 | 0.706 | 0.761 | 1.000 | 20816.3 | 11553.7 |
| XBB.1.16.9 | Canada | 0.732 | 0.673 | 0.786 | 1.001 | 19067.4 | 11645.8 |
| DV.7.1.1 | Canada | 0.732 | 0.689 | 0.772 | 1.000 | 22018.8 | 11793.1 |
| EG.6.1.1 | Canada | 0.711 | 0.653 | 0.764 | 1.000 | 20595.7 | 12291.5 |
| GE.1 | Canada | 0.690 | 0.624 | 0.751 | 1.000 | 23013.5 | 11109.6 |

**Table S4. Primers used in this study**

| Primer name | Primer sequence (5'-to-3') | Purpose |
|---|---|---|
| Omicron universal Fw | cactatagggcgaattgggtaccatgtttgtgttcctggt | Preparation of S expression plasmid |
| BA.2 WT Rv | agctccaccgcggtggcggccgctcaggtgtagtgcagtttca | Preparation of S expression plasmid |
| PJ4601 R346T Fw | gtgttcaatgccaccACCtttgcctctgtctat | Preparation of S expression plasmid |
| PJ4602 R346T Rv | atagacagaggcaaaGGTggtggcattgaacac | Preparation of S expression plasmid |
| PJ4603 F456L Fw | tacTGGtacagaagcCTGaggaagagcAAGctg | Preparation of S expression plasmid |
| PJ4604 F456L Rv | cagCTTgctcttcctCAGgcttctgtaCCAgta | Preparation of S expression plasmid |
| PJ4605 V1104L Fw | ggcacccactggtttCTGacccagaggaacttc | Preparation of S expression plasmid |
| PJ4605 V1104L Rv | gaagttcctctgggtCAGaaaccagtgggtgcc | Preparation of S expression plasmid |

### Supplementary figure

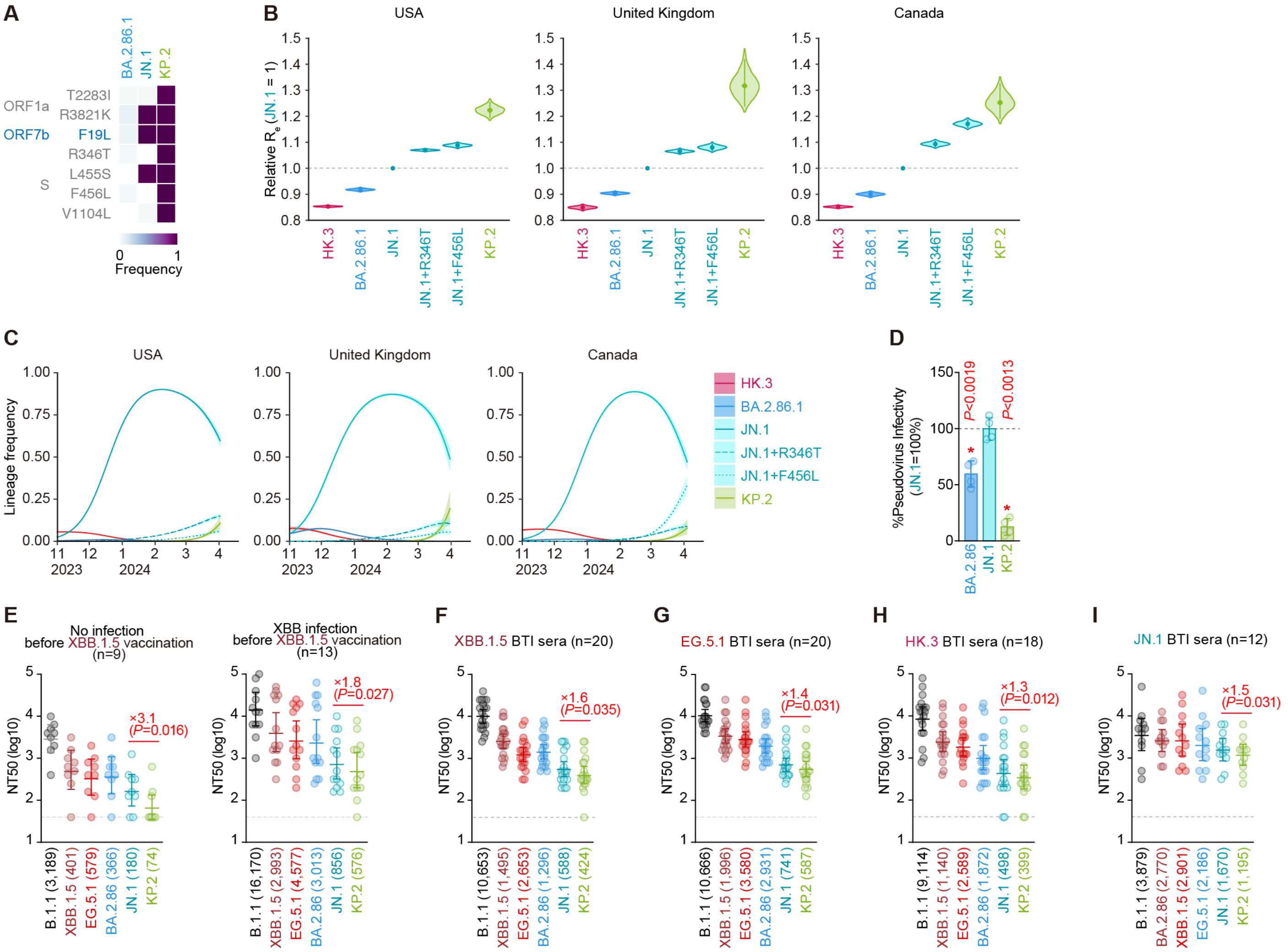


**Figure S1. Virological features of KP.2**

(**A**) Frequency of mutations in KP.2 and other lineages of interest. Only mutations with a frequency >0.5 in at least one but not all the representative lineages are shown.

(**B**) Estimated relative $R_e$ of the variants of interest in the USA, United Kingdom, and Canada. The relative $R_e$ of JN.1 is set to 1 (horizontal dashed line). Violin, posterior distribution; dot, posterior mean; line, 95% Bayesian confidence interval.

(**C**) Estimated epidemic dynamics of the variants of interest in the USA, United Kingdom, and Canada from November 1, 2023 to April 11, 2024. Countries are ordered according to the number of detected sequences of KP.2 from high to low. Line, posterior mean, ribbon, 95% Bayesian confidence interval.

(**D**) Lentivirus-based pseudovirus assay. HOS-ACE2/TMPRSS2 cells were infected with pseudoviruses bearing each S protein of BA.2.86, JN.1 and KP.2. The amount of input virus was normalized to the amount of HIV-1 p24 capsid protein. The percentage infectivity of BA.2.86 and KP.2 are compared to that of JN.1. The horizontal dash line indicates the mean value of the percentage infectivity of JN.1. Assays were performed in quadruplicate, and a representative result of four independent assays is shown. The presented data are expressed as the average ± SD. Each dot indicates the result of an individual replicate. Statistically significant differences versus JN.1 is determined by two-sided Student's *t* tests (*, $P<0.01$).

(**E**-**I**) Neutralization assay. Assays were performed with pseudoviruses harboring the S proteins of B.1.1, XBB.1.5, EG.5.1, BA.2.86, JN.1 and KP.2. The following sera were used: vaccinated sera from fully vaccinated individuals who had not been infected (9 donors) and vaccinated sera from fully vaccinated individuals who had been infected with XBB subvariants (after June, 2023) (13 donors) (**E**); convalescent sera from fully vaccinated individuals who had been infected with XBB.1.5 (one 2-dose vaccinated donor, seven 3-dose vaccinated donors, six 4-dose vaccinated donors, five 5-dose vaccinated donors and one 6-dose vaccinated donor. 20 donors in total) (**F**); EG.5.1 (one 2-dose vaccinated donor, six 3-dose vaccinated donors, five 4-dose vaccinated donors, four 5-dose vaccinated donors and four 6-dose vaccinated donors. 20 donors in total) (**G**); individuals who had been infected with HK.3 (three 2-dose vaccinated donors, five 3-dose vaccinated donor, two 4-dose vaccinated donors, three 5-dose vaccinated donors, one 6-dose vaccinated donor and four donors with unknown vaccine history. 18 donors in total) (**H**) and individuals who had been infected with JN.1 (one 2-dose vaccinated donor, two 3-dose vaccinated donors, two 7-dose vaccinated donors and seven donors with unknown vaccine history. 12 donors in total) (**I**). Assays for each serum sample were performed in quadruplicate to determine the 50% neutralization titer ($NT_{50}$). Each dot represents one $NT_{50}$ value, and the geometric mean and 95% confidence interval are shown. The number in parenthesis indicates the geometric mean of $NT_{50}$ values. The horizontal dash line indicates the detection limit (40-fold). Statistically significant differences versus KP.2 were determined by two-sided Wilcoxon signed-rank tests, and p values are indicated in parentheses. The fold changes of $NT_{50}$ from that of KP.2 are indicated with "X".

**Consortia**

**The Genotype to Phenotype Japan (G2P-Japan) Consortium**

**The Institute of Medical Science, The University of Tokyo, Japan**
Naoko Misawa, Arnon Plianchaisuk, Ziyi Guo, Alfredo Hinay Jr., Kaoru Usui, Wilaiporn Saikruang, Spyridon Lytras, Ryo Yoshimura, Shusuke Kawakubo, Luca Nishimura, Shigeru Fujita, Luo Chen, Jarel Elgin M. Tolentino, Lin Pan, Wenye Li, Maximilian Stanley Yo, Kio Horinaka, Mai Suganami, Adam P. Strange, Mika Chiba, Keiko Iida, Naomi Ohsumi, Shiho Tanaka, Eiko Ogawa, Kyoko Yasuda, Tsuki Fukuda, Rina Osujo

**Hokkaido University, Japan**
Takasuke Fukuhara, Tomokazu Tamura, Rigel Suzuki, Saori Suzuki, Hayato Ito, Keita Matsuno, Hirofumi Sawa, Naganori Nao, Shinya Tanaka, Masumi Tsuda, Lei Wang, Yoshikata Oda, Zannatul Ferdous, Kenji Shishido, Keita Mizuma, Isshu Kojima, Jingshu Li, Tomoya Tsubo, Shuhei Tsujino

**Tokai University, Japan**
So Nakagawa

**Kyoto University, Japan**
Kotaro Shirakawa, Akifumi Takaori-Kondo, Kayoko Nagata, Ryosuke Nomura, Yoshihito Horisawa, Yusuke Tashiro, Yugo Kawai, Kazuo Takayama, Rina Hashimoto, Sayaka Deguchi, Yukio Watanabe, Ayaka Sakamoto, Naoko Yasuhara, Takao Hashiguchi, Tateki Suzuki, Kanako Kimura, Jiei Sasaki, Yukari Nakajima, Hisano Yajima, Yoshitaka Nakata, Hiroki Futatsusako

**Hiroshima University, Japan**
Takashi Irie, Ryoko Kawabata

**Kyushu University, Japan**
Kaori Tabata

**Kumamoto University, Japan**
Terumasa Ikeda, Hesham Nasser, Ryo Shimizu, MST Monira Begum, Michael Jonathan, Yuka Mugita, Otowa Takahashi, Kimiko Ichihara, Takamasa Ueno, Chihiro Motozono, Mako Toyoda, Sharee Leong

**University of Miyazaki, Japan**
Akatsuki Saito, Maya Shofa, Yuki Shibatani, Tomoko Nishiuchi

**Tokyo Metropolitan Institute of Public Health, Japan**
Kazuhisa Yoshimura, Kenji Sadamasu, Mami Nagashima, Hiroyuki Asakura, Isao Yoshida

**Charles University, Czechia**
Prokopios Andrikopoulos, Miguel Padilla-Blanco, Aditi Konar

**Acknowledgments**

We would like to thank all members of The Genotype to Phenotype Japan (G2P-Japan) Consortium. We thank Kenzo Tokunaga (National Institute of Infectious Diseases, Japan) for sharing materials and Yuka Kamoshita (Department of Laboratory Medicine, Keio University School of Medicine) and Masayo Noguchi (Clinical Laboratory, Keio University Hospital) for supporting patient sera collection. We gratefully acknowledge the numerous laboratories worldwide that have provided sequence data and metadata to GISAID. A full list of originating and submitting laboratories for the sequences used in our analysis can be found at https://www.gisaid.org using the EPI-SET-ID: EPI-SET-ID: EPI_SET_240418tb.
